# Long-lasting germinal center responses to a priming immunization with continuous proliferation and somatic mutation

**DOI:** 10.1101/2021.12.20.473537

**Authors:** Jeong Hyun Lee, Henry Sutton, Christopher A. Cottrell, Ivy Phung, Gabriel Ozorowski, Leigh M. Sewall, Rebecca Nedellec, Catherine Nakao, Murillo Silva, Sara T. Richey, Jonathan L. Torres, Wen-Hsin Lee, Erik Georgeson, Michael Kubitz, Sam Hodges, Tina-Marie Mullen, Yumiko Adachi, Kimberly M. Cirelli, Amitinder Kaur, Carolina Allers-Hernandez, Marissa Fahlberg, Brooke F. Grasperge, Jason P. Dufour, Faith Schiro, Pyone P. Aye, Diane G. Carnathan, Guido Silvestri, Xiaoying Shen, David C. Montefiori, Ronald S. Veazey, Andrew B. Ward, Lars Hangartner, Dennis R. Burton, Darrell J. Irvine, William R. Schief, Shane Crotty

**Affiliations:** International AIDS Vaccine Initiative Neutralizing Antibody Center, La Jolla 92037, CA 92037, USA; Center for Infectious Disease and Vaccine Research, La Jolla Institute for Immunology (LJI), La Jolla, CA 92037, USA; Biomedical Sciences (BMS) Graduate Program. School of Medicine, University of California, San Diego (UCSD), La Jolla, CA 92037, USA; Consortium for HIV/AIDS Vaccine Development (CHAVD), The Scripps Research Institute, La Jolla, CA 92037 USA; Koch Institute for Integrative Cancer Research, Massachusetts Institute of Technology, Cambridge, MA 02139 USA; Ragon Institute of Massachusetts General Hospital, Massachusetts Institute of Technology and Harvard University, Cambridge, MA 02139 USA; Department of Biological Engineering, Massachusetts Institute of Technology, Cambridge, MA 02139 USA; Department of Immunology and Microbiology, The Scripps Research Institute, La Jolla, CA 92037 USA; Department of Integrative Structural and Computational Biology, The Scripps Research Institute, La Jolla, CA 92037 USA; IAVI Neutralizing Antibody Center, The Scripps Research Institute, La Jolla, CA 92037 USA; Tulane National Primate Research Center, Tulane School of Medicine, Covington, LA 70433; Yerkes National Primate Research Center and Emory Vaccine Center, Emory University School of Medicine, Atlanta, GA 30322; Department of Surgery, Laboratory for AIDS Vaccine Research & Development, Duke University Medical Center, Duke University, Durham, NC 27710; Department of Materials Science and Engineering, Massachusetts Institute of Technology, Cambridge, MA 02139 USA; Howard Hughes Medical Institute, Chevy Chase, MD 20815 USA; Department of Medicine, Division of Infectious Diseases and Global Public Health, University of California, San Diego (UCSD), La Jolla, CA 92037, USA

## Abstract

Germinal centers (GCs) are the engines of antibody evolution. Using HIV Env protein immunogen priming in rhesus monkeys (RM) followed by a long period without further immunization, we demonstrate GC B cells (B_GC_) lasted at least 6 months (29 weeks), all the while maintaining rapid proliferation. A 186-fold B_GC_ cell increase was present by week 10 compared to a conventional immunization. Single cell transcriptional profiling revealed that both light zone and dark zone GC states were sustained throughout the 6 months. Antibody somatic hypermutation (SHM) of B_GC_ cells continued to accumulate throughout the 29 week priming period, with evidence of selective pressure. Additionally, Env-binding B_GC_ cells were still 49-fold above baseline 29 weeks after immunization, suggesting that they could be active for significantly longer periods of time. High titers of HIV neutralizing antibodies were generated after a single booster immunization. Fully glycosylated HIV trimer protein is a complex antigen, posing significant immunodominance challenges for B cells, among other difficulties. Memory B cells (B_Mem_) generated under these long priming conditions had higher levels of SHM, and both B_Mem_ cells and antibodies were more likely to recognize non-immunodominant epitopes. Numerous B_GC_ cell lineage phylogenies spanning the >6-month GC period were identified, demonstrating continuous GC activity and selection for at least 191 days, with no additional antigen exposure. A long prime, adjuvanted, slow delivery (12-day) immunization approach holds promise for difficult vaccine targets, and suggests that patience can have great value for tuning GCs to maximize antibody responses.

## Main

Antibodies serve as effective adaptive immunity frontline defenses against most infectious diseases. As such, most efficacious vaccines aim to prophylactically elicit potent neutralizing antibodies and long-lasting immunological memory to the target pathogen. For rapidly mutating pathogens such as HIV, there is an additional level of complication wherein an ideal vaccine should generate cross-reactive or broadly neutralizing antibodies (bnAbs) that can protect against variants^1,2^, but to date bnAbs against HIV have not yet been elicited in humans nor non-human primates (NHPs) by vaccination^3^.

High affinity antibodies are typically the result of affinity maturation through evolutionary competition among B cells in GCs. GCs are evolution in miniature, with proliferation (generations) accompanied by mutations, and competition for limiting resources in the form of antigen and T cell help^4–7^. To accomplish this evolution, B_GC_ cells proliferate rapidly — every 4-6 hours^8,9^. GCs are often observed for a few weeks after an acute antigen exposure. Antigen-specific B_GC_ cells have widely been observed for 14 to 28 days in most model systems, and such a time window can represent a substantial amount of antibody sequence space exploration by B_GC_ due to their fast cell cycle^4,7^. We previously showed that vaccine slow delivery methods over a period of 7 to 14 days, such as the use of osmotic pumps or repeated small dose injections, enhanced GC responses relative to traditional bolus immunizations in terms of magnitude of B_GC_ cells and antibody responses^10,11^, with some evidence of increasing the durability of GCs for two months^12^. However, the full potential longevity of GCs, the biological programming of older GCs, antibody maturation under such conditions, and the functionality and productivity of older GCs are minimally understood. Here, we utilized a 12-day slow delivery protein immunization strategy and antigen-specific molecular and cellular tools to explore the extent of GC durability after a priming immunization and the ensuing immunological outcomes.

### A priming immunization can fuel GCs for months

Alum is a classic adjuvant used in many human vaccines^13^. A group of rhesus monkeys (RMs) were given bolus injections of recombinantly expressed stabilized HIV Env trimer^14^, MD39 (50 µg protein) formulated with alum adjuvant (Alhydrogel), reflective of how most licensed human protein vaccines are formulated and administered (Group 1, Fig. 1a). In an effort to generate more robust GCs, we immunized two groups of RMs with MD39 Env trimer formulated with the new ISCOM-type adjuvant saponin/MPLA nanoparticle (SMNP)^15^ (Groups 2 & 3, Fig. 1a). The priming immunization for these two groups was administered via a slow delivery vaccination method termed escalating dose (ED)^10^, where the total dose of the MD39 plus SMNP formulation (50 µg protein, 375 µg adjuvant per side) was split between 7 gradually increasing doses, delivered every other day for a total of 12 days (Extended Data Fig. 1a). Group 3 was designed with an unusual “long prime” period to assess the durability of GCs after a primary immunization. Each animal in the study was immunized bilaterally, thereby doubling the number of lymph nodes (LNs) and GCs that could be tracked over time. GCs were sampled every 2-3 weeks by LN fine needle aspiration (FNA) of inguinal LNs (ILNs) (Fig. 1a).

**Fig. 1:**
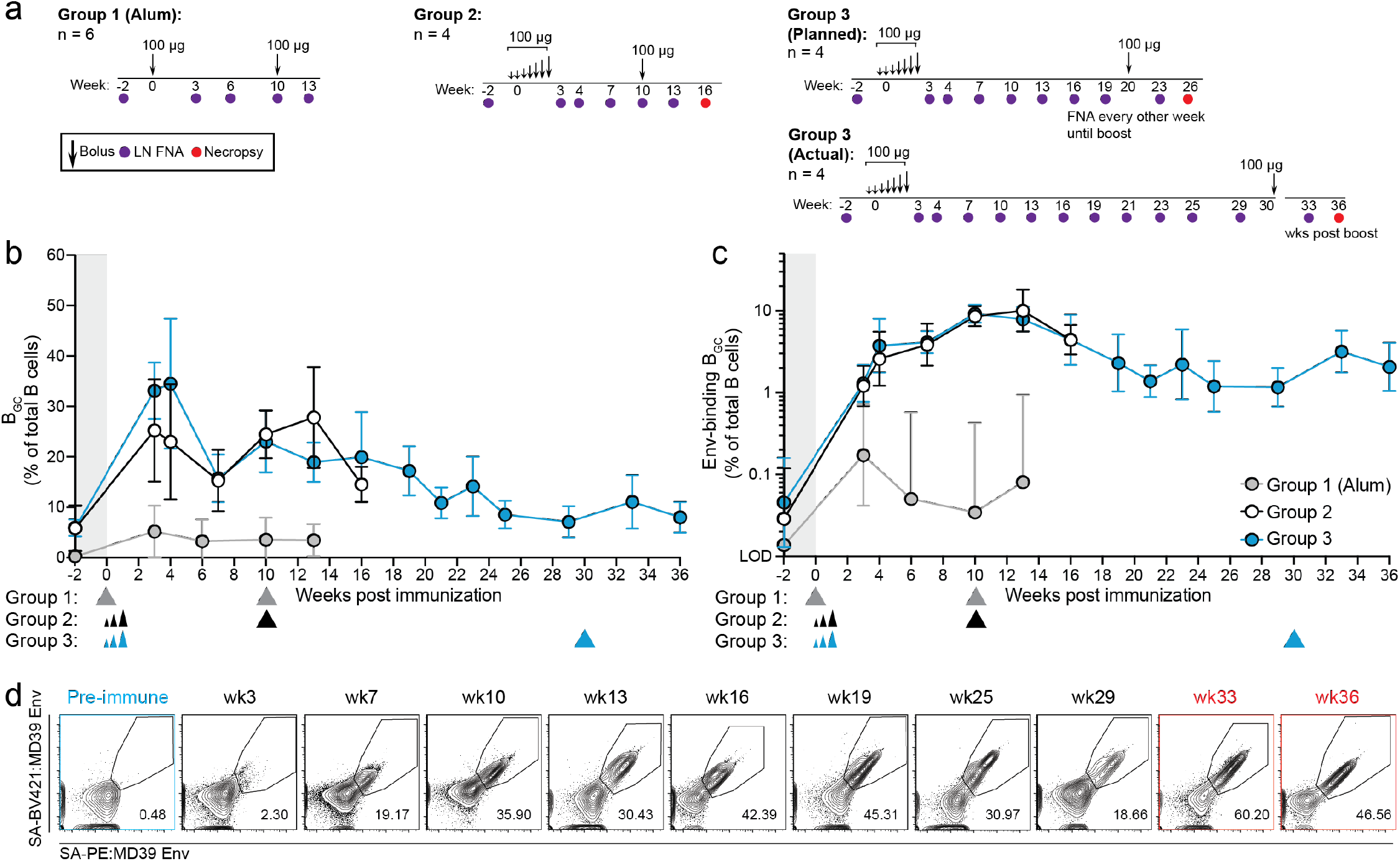
GCs following a priming immunization can last for over half a year. **a**, Experimental schematic. **b-c**, Quantification of longitudinal BGC cell kinetics. Left and right ILN FNA samples are independent data points. Triangles below indicate the prime and boost immunization time points. **b**, Quantification of total BGC cells as a percent of total CD20^+^ B cells. **c**, Env-binding BGC cells as a percent of total B cells. Left and right ILNs are graphed as independent data points. **d**, Longitudinal detection of Env-binding BGC cells in the “Long Prime” Group 3. Representative FACS plots from left ILN of one animal. Red indicates post-boost. Gated on CD20^+^/CD71^+^CD38^-^ BGC cells. Mean and SD or geometric mean and geometric SD are plotted depending on the scale. Limit of detection (LOD). Mann-Whitney test: *P < 0.05; **P < 0.005.

Following the priming immunization, conventional MD39 plus alum—bolus immunized animals (Group 1) exhibited an increase in total B_GC_ cell (CD71^+^CD38^-^) % at week 3 post-immunization (Fig. 1b); the frequency of Env-binding B_GC_ cells (CD71^+^CD38^-^/Env^+/+^) peaked at week 3 and declined thereafter (Env-binding B_GC_ cells as % of total B cells. Fig. 1c). Both total and Env-binding B_GC_ cells were substantially increased in MD39 plus SMNP ED—immunized RMs compared to RMs that received conventional protein plus alum bolus immunization (Groups 2 & 3 combined vs. Group 1, Fig. 1b-c, Extended Data Fig. 1b, 2a). Median peak B_GC_ cell frequencies observed were 24-33% compared to 3.5% (Mann-Whitney P < 0.0001, week 3 Groups 2 & 3 combined vs. Group 1. Fig. 1b). Median Env-binding B_GC_ cell frequencies were approximately 7.8-times greater at week 3 (Mann-Whitney, P < 0.0025, Groups 2 & 3 combined vs Group 1. Fig. 1c). Strikingly, in contrast to the conventionally primed Group 1, frequencies of Env-binding B_GC_ cells in Groups 2 and 3 continued to increase, resulting in a 186-fold GC difference by week 10 (Mann-Whitney, P < 0.0001, Group 2 & 3 combined vs Group 1. Fig. 1c)

Tracking of the priming immune response continued for Group 3 animals beyond week 10 (Group 1 & 2 animals were boosted at week 10, Fig. 1a). Surprisingly, GC responses were still active at weeks 13, 16, 21, 25, and 29 (Fig. 1b-d, Extended Data Fig. 2a-b). The median magnitude of these Env-binding B_GC_ cells at week 29 was still 27-fold higher than the peak Env-binding B_GC_ cells observed after conventional alum immunization, and it was also greater than the post-boost Env-binding B_GC_ cell response to conventional alum immunization (Fig. 1c). 191 days (27 weeks) after the end of the priming dose (29 weeks from day 0), median Env-binding B_GC_ cell frequencies in ILNs were still ∼49-fold higher than baseline (Fig. 1b-d, Extended Data Fig. 2a-b). Thus, GCs were capable of continuous activity for > 191 days with no additional antigen introduced.

GC-T follicular helper (T_FH_) cells play a critical role in the recruitment and selection of B_GC_ cells^5^. Total GC-T_FH_ cell frequencies in ILNs changed over the course of the priming period (Extended Data Fig. 2c), however longitudinal quantitation of Env-specific GC-T_FH_ cells was not possible due to limiting FNA samples. At 6 weeks post-boost, increased Env-specific GC-T_FH_ cell frequencies trended higher in the long prime Group 3 (Extended Data Fig. 2d-e)^16^. Long-lasting prime GCs may contribute to an improved antigen-specific GC-T_FH_ response after the booster immunization.

### Enhanced antibody response quality

Group 3 RMs Env-binding serum IgG titers remained stable from week 3 to 29 of the priming phase in the absence of a booster immunization (Fig. 2a). After boosting, Group 2 and 3 animals generated similar peak binding antibody titers (2 to 3 weeks post-boost), but Group 3 animals maintained significantly higher Env-binding IgG titers at week 6 post-boost (Fig. 2b). The quality of the antibody responses was next evaluated in terms of ability to neutralize the tier-2 autologous BG505 pseudovirus. Notably, autologous tier-2 neutralizing antibodies were detectable in all long prime Group 3 animals after only the priming immunization (geometric mean titer [GMT] ∼170 at week 29, Fig. 2c). All animals receiving ED immunization generated robust neutralizing antibody responses post-boost (Fig. 2c-d); in contrast, only a single animal with conventional bolus immunization adjuvanted with alum had detectable autologous neutralizing antibodies, which were of low titer (∼37, Extended Data Fig. 3a). Peak observed autologous tier-2 neutralizing antibody GMTs in Group 3 (long prime) were all > 2,000 (week 3 post-boost, Fig. 2c). Group 3 autologous tier 2 neutralization titers were 4-fold greater than Group 2 at 6 weeks post-boost (Fig. 2c-d). The titers described represent the most robust and consistent autologous tier-2 neutralizing antibody responses in RMs after two immunizations in any of our studies^11,12,15^.

**Fig 2:**
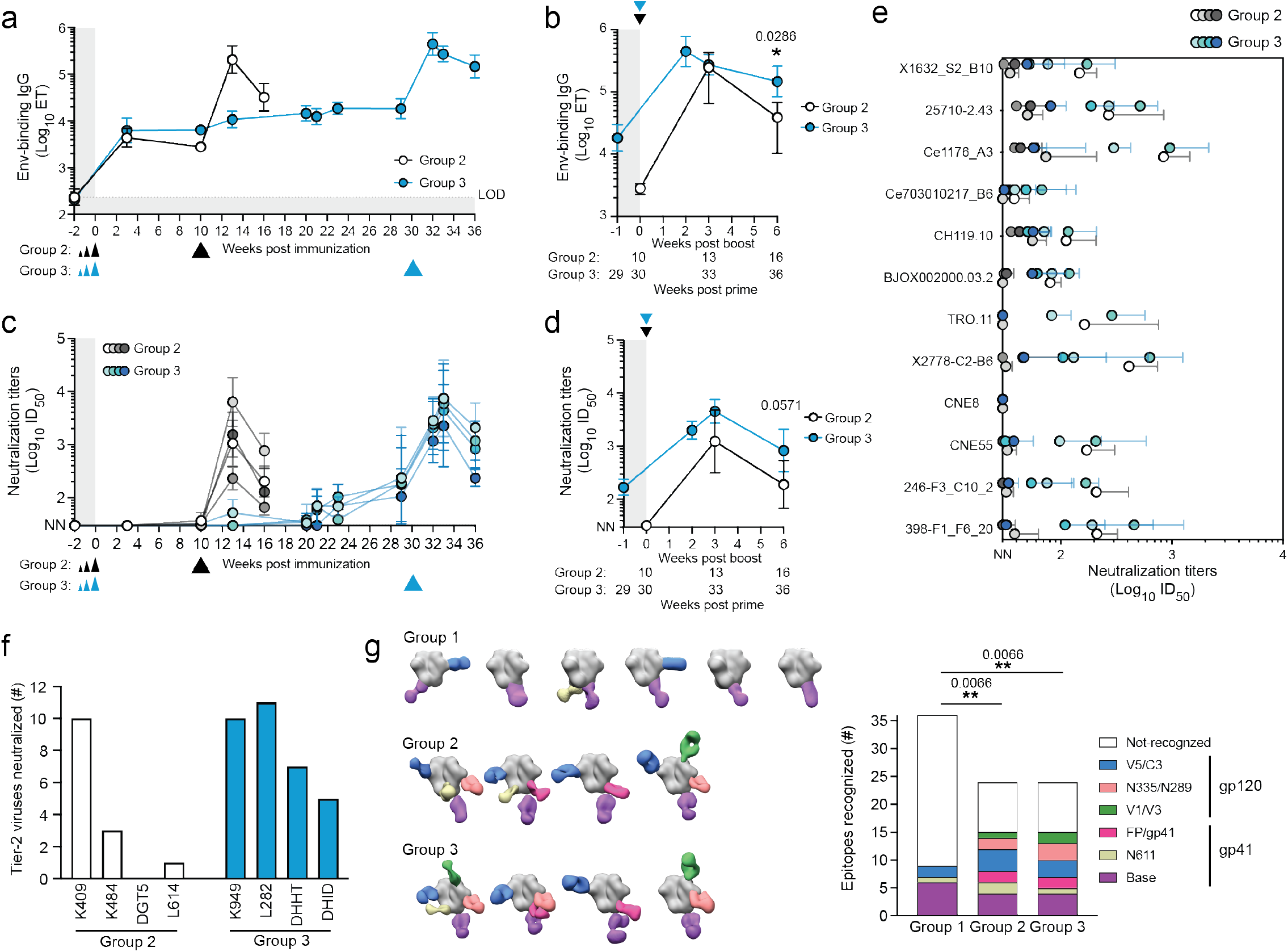
Long priming enhanced antibody quality. **a**, Env-binding serum IgG titers determined by ELISA. ET: Endpoint titer. **b**, Env-binding IgG titers following boost. Triangles indicate the boost time point. **c**, BG505 pseudovirus neutralization titers at 50 % inhibition (ID50). **d**, BG505 neutralization titers at post-boost time points. **e**, Heterologous tier-2 virus neutralization titers. ID50 ≤ 30 was considered non-neutralizing (NN). **f**, Number of tier-2 heterologous viruses neutralized (ID50 > 50) in the 12-virus panel by week 3 post-boost serum. **g**, EMPEM of polyclonal plasma Fabs post-boost. Group 1 at week 2 and Groups 2 & 3 at week 3 post-boost. The Env trimer is shown in gray. Graphs quantify number of animals recognizing each indicated epitope. The color of the Fabs in the EM map match that of the epitope colors in the bar graph. Fisher’s exact test comparing number of epitopes recognized vs. not recognized between groups, **P < 0.01.

Groups 2 and 3 post-boost sera exhibited some of the broadest observed tier-2 neutralizing antibody specificities elicited by Env trimer immunization, with most animals from Group 3 exhibiting greater breadth than those in Group 2 (Fig. 2e-f). The 12-virus neutralization panel was repeated by an independent laboratory, with similar observation of neutralization breadth (Extended Data Fig. 3b-c). In head-to-head neutralization assays, limited tier-2 neutralization breadth was observed with serum from an earlier RM study given ED immunization and two booster immunizations with a similar Env trimer (Olio6) and an earlier ISCOMs adjuvant (SMNP without MPLA)^12^ (Extended Data Fig. 3d-h), and likewise for serum from an RM study with three bolus immunizations of MD39 Env trimer and SMNP (Extended Data Fig. 3e-h)^15^.

The vast majority of HIV Env-binding B cells and antibodies are usually directed to immunodominant non-neutralizing epitopes, such as the base of soluble recombinant Env trimers^12,17–20^. Electron microscopy polyclonal epitope mapping (EMPEM)^19^ of circulating antibodies revealed that the number of targeted epitopes correlated with autologous neutralizing antibody titers (Fig. 2g and Extended Data Fig. 4). Group 2 and 3 animals generated antibody responses to V5/C3 and V1/V3 epitopes associated with autologous BG505 SHIV protection^21^. Antibody responses in conventional bolus plus alum-immunized animals were largely restricted to the Env trimer base (Fig. 2g). In sum, employing a 12-day ED immunization strategy and vaccine formulation with SMNP was associated with substantially improved epitope breadth and quality of neutralizing antibodies.

### Six-month B_GC_ cells are highly functional

B_GC_ cell characteristics after priming were subsequently interrogated in greater detail to assess their functionality over time, given the apparent presence of continuously active B_GC_ cells for over six months. BCL6 is the lineage-defining transcription factor of B_GC_ cells and is essential for their functionality^6^. KI67 (*MKI67*) marks rapidly dividing cells. LN B cells from month 5 to 6 (week 21-25) were stained for BCL6 and KI67 protein. On average, ∼72% of Env-binding CD71^+^CD38^-^ B_GC_ cells were KI67^+^BCL6^+^ (Fig. 3a-b and Extended Data Fig. 5), indicating retained B_GC_ programming and proliferation for at least six months. To further ascertain the phenotypic and functional characteristics of B_GC_ cells at different time points over the course of six months, single cell transcriptional profiling was done for ∼70,000 cells from LN FNAs of weeks 3, 4, 7, 10, 13, 16, 29, and 33, predominantly consisting of Env-binding B_GC_ cells, as well as peripheral blood sorted Env-binding memory B cells (B_Mem_) from weeks 16 (Group 2) and 36 (Group 3). Dark zone (DZ) and light zone (LZ) cell clusters were clearly observed among LN B cells when analyzing all time points together (Fig. 3c-d, Extended Data Fig. 6). We then examined the B_GC_ cell transcriptional profiles over the course of the Group 3 RMs long priming period (week 3, 7, 16, 29. Fig. 3d-h, Extended Data Fig. 6)^22,23^. LZ and DZ states were sustained across the six-month period (Fig. 3d). Expression of key functional B_GC_ genes *MKI67, AICDA, MYC*, and *CD40* were maintained over time, and were compartmentalized comparably between DZ and LZ cell types at all time points (Fig. 3e-h). The ratio of DZ:LZ cells remained relatively consistent over the course of the priming period (Fig. 3i). Overall, antigen-specific B_GC_ cells possessed stable phenotypic characteristics over a six-month period, indicative of long-term maintenance of functional B_GC_ cell properties in the absence of additional immunization.

**Fig. 3:**
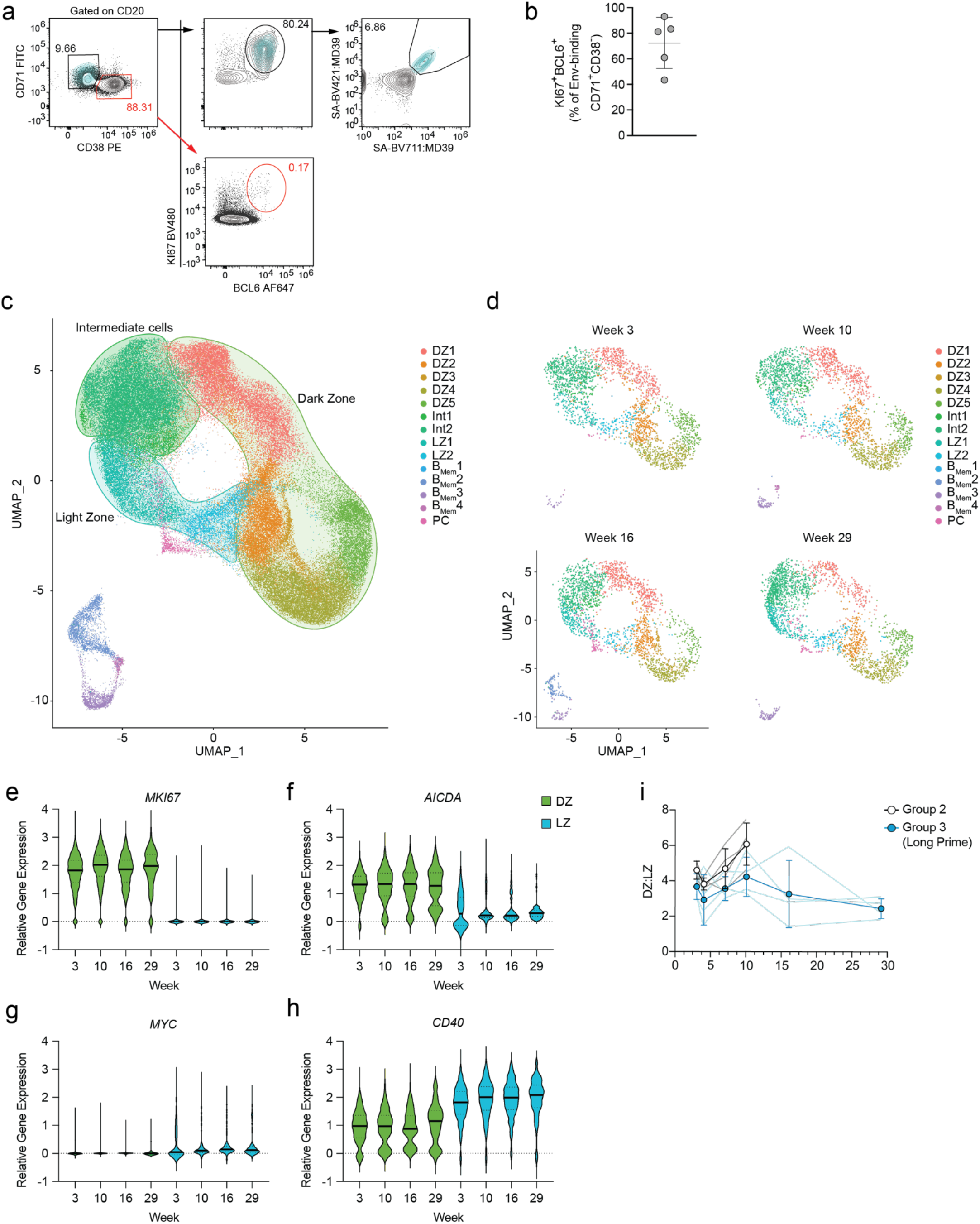
BGC cell phenotypic and functional characteristics over the course of six months. **a**, Representative flow cytometry gating showing CD71^+^CD38^-^ and Ki67^+^BCL6^+^ BGC cells. Back-gating of CD71^+^CD38^-^/Ki67^+^BCL6^+^/Env^+/+^ BGC cells is shown in cyan. CD71^-^CD38^+^ non-BGC cells are not Ki67^+^BCL6^+^ (red gates). **b**, Frequency of KI67^+^BCL6^+^ cells among CD71^+^CD38^-^/Env^+/+^ BGC. **c**, UMAP projection of single cell gene expression profiles identifying clusters of B cell states from LN FNA and PBMCs. Int, LZ-DZ Intermediate populations. PC, Plasma cells. **d**, Per time point UMAP plots extracted from (d). **e-h**, relative gene expression of *MKI67* (f), *AICDA* (g), *MYC* (h), *CD40* (i) in the DZ (DZp3) and LZ (LZ2). **i**, DZ:LZ ratio as determine by single cell clustering in the LN after priming.

### B_GC_ cell BCR evolution for months in the absence of additional antigen

To directly assess functionality of the GCs over these extended time periods, multiple experimental approaches were employed comparing Groups 2 and 3 RMs. We performed BCR sequencing of Env-binding LN FNA-derived B_GC_ cells from nine different timepoints to assess SHM over time, as well as clonal diversity and mutational patterns in clonal lineages (Fig. 4 and Extended Data Fig. 7-10). Env-binding B_GC_ cell heavy chain (HC) nucleotide (NT) mutations increased significantly between week 3 and 10 (Mann-Whitney, P < 0.0001 for both Group 2 & 3, Fig. 4a-b). Notably, B_GC_ cells continued to accumulate mutations in the absence of another immunization through week 29 in Group 3 RMs, at which point the median number of HC mutations was 17, with the top 25% of B_GC_ cells containing 22 to 45 HC mutations (Fig. 4a-b). The difference in SHM in the long prime (week 29) versus 10-week prime was highly significant (Mann-Whitney, P < 0.0001, Fig. 4b; P < 0.0009, Fig. 4c), and the difference in median mutations between weeks 10 and 29 was nearly as great as the difference between weeks 3 and 10, indicative of robust GC functionality continuing through at least week 29 (Fig. 4a-c). Env-binding B_GC_ cells showed a gradual reduction in the diversity of clones (population diversity) over time (Fig. 4d-e), further indicative of ongoing competitive pressure. The proportion of unmutated Env-binding B_GC_ cells dropped over time, with 0.19-0.42% unmutated cells by week 7 (Fig. 4b). Substantial mutations were also observed in the light chain (LC) sequences over time, with comparable patterns to the HCs (Extended Data Fig. 7a-b).

**Fig. 4:**
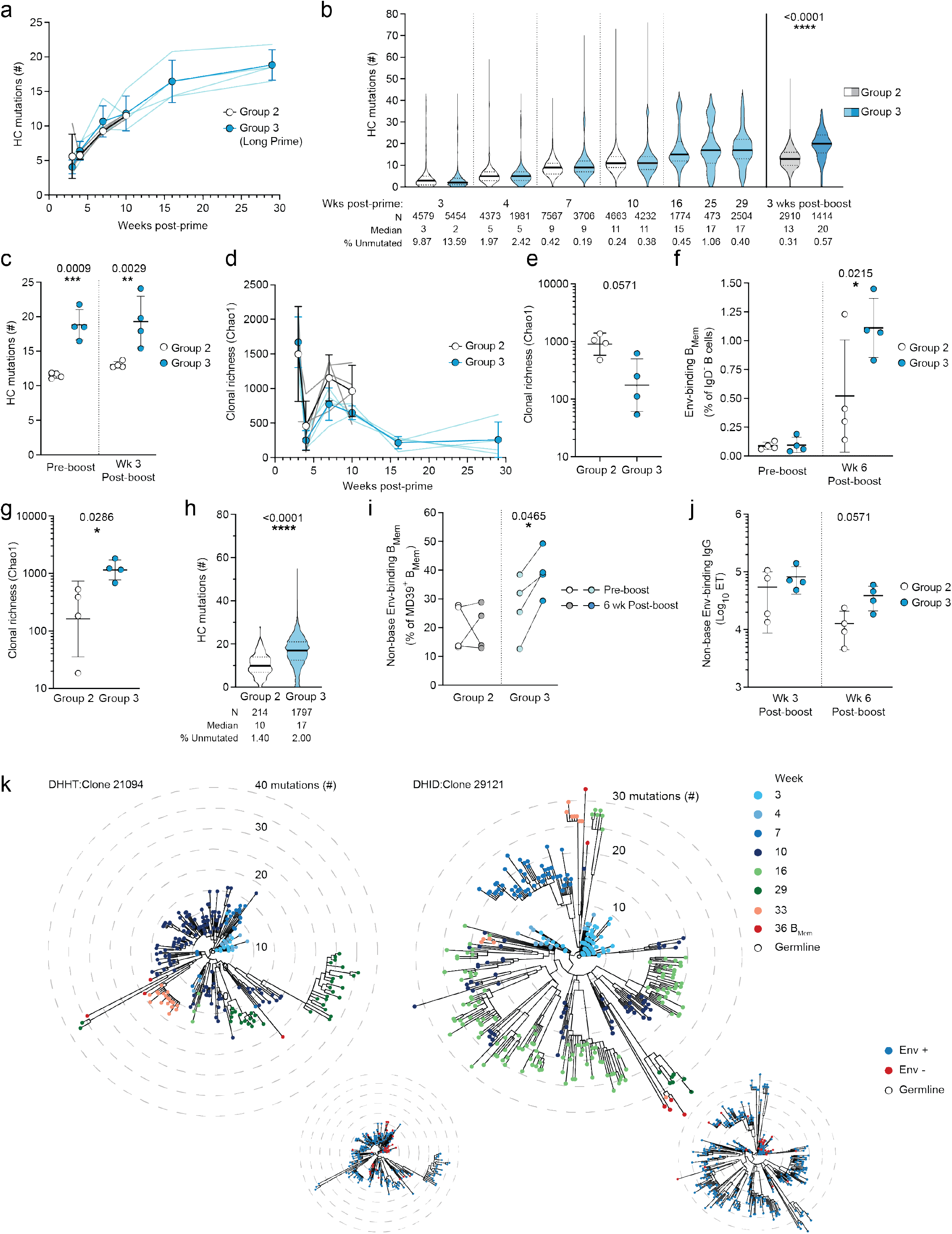
Clonal competition and affinity maturation occurs in antigen-specific BGC cells identified in long lasting GCs. **a**, Number of nucleotide (NT) mutations in the HC (VH + JH) of Env-binding BGC cells after priming, spaghetti plots track mutations per animal. **b**, Number of pre- and post-boost NT mutations in Env-binding BGC cells. **c**, Comparison of BGC mutations at the last pre-boost time point (pre-boost; week 10 and 29 for Group 2 & 3 respectively) and 3 weeks post-boost (week 13 and 33 for Group 2 & 3 respectively). 2-way ANOVA multiple comparisons test. **d**, BGC population diversity at post-prime time points (Chao1). **e**, BGC cell pre-boost population diversity. pre-boost; week 10 and 29 for Group 2 & 3 respectively. **f**, Frequency of Env-binding BMem cells in blood. 2-Way ANOVA multiple comparisons test. **g**, Clonal diversity of BMem cells after boosting. **h**, The number of mutations in week 6 post-boost BMem cells. **i**, Quantitation of Env-binding BMem cells that do not bind the trimer-base epitope. **j**, Serum titers of non-base directed Env-binding antibodies detected by ELISA. **k**, Clonal trees of 2 unique B cell lineages from two different long prime Group 3 animals. The tree on the left is color coded by time points while the tree on the right is color coded by Env binding. Each ring indicates 5 HC mutations from the predicted germline. For all graphs, Mann-Whitney test was used unless otherwise indicated. ns > 0.05, *P < 0.05, **P < 0.01, ***P < 0.001, ****P < 0.0001.

After the booster immunization, HC mutations increased in Env-binding B_GC_ cells of both Group 2 and 3 RMs, with the highest overall number of mutations in the long primed RMs (week 3 post-boost HC median mutations = 13 vs. 20, Fig. 4b-c, Extended Data Fig. 7c). Comparable observations were made for LC mutations (Extended Data Fig. 7d).

Pre-boost Env-binding B_Mem_ cell (CD20^+^IgD^—^Env^+/+^) frequencies in peripheral blood were equivalent in RMs from Groups 2 and 3 (Fig. 4f, Extended Data Fig. 7e). Boosting increased Env-binding B_Mem_ frequencies in both groups (Fig. 4f). RMs with the long prime had more highly mutated B_Mem_ cells and greater clonal richness among B_Mem_ cells (Fig. 4g-h, Extended Data Fig. 7f). This was also reflected in a significant shift away from immunodominant base-binding Env-specific B_Mem_ cells of the long primed group compared to Group 2 (Fig. 4i), a phenomenon that was also reflected in the circulating antibody titers (Fig. 4j).

Clonal lineage analysis of paired BCR sequences was used to examine GC duration and functionality over months after a single immunization. Many clonal lineages were identified with B_GC_ clones observed at the first and last LN FNA timepoint post-prime, with extensive SHMs in-between (Fig. 4k, Extended Data Fig. 8-10). Some clonal lineages were identified with B_GC_ clones observed at every, or nearly every, LN FNA timepoint, providing direct evidence of B_GC_ cell persistence for 29 weeks (Fig. 4k, Extended Data Fig. 8, 9a, 10a-b). Diversification of clones in clonal lineages was apparent. Daughter clones most evolutionarily divergent from the germline were typically present at late time points (mutations to time correlation. R^2^ = 0.72, P < 0.001, lineage 21094; R^2^ = 0.68, P < 0.001, lineage 29121. Fig. 4k, Extended Data Fig. 8, 9d-e). For some lineages such as 5491 and 29183, the majority of early B_GC_ cell clones did not bind Env by flow cytometry (Env^—/—^), but the majority of B_GC_ clones from week 16+, and B_Mem_ clones from week 36, bound Env by flow cytometry (Env^+/+^), indicative of a clone that started with low affinity to Env and affinity matured (Extended Data Fig. 10a).

B_Mem_ cells were commonly represented in the clonal lineage trees among multiple sub-lineages, including amongst the most mutated branches observed at late LN FNA B_GC_ time points (Fig. 4k; Extended Data Fig. 8-10), demonstrating that the GCs productively output B_Mem_ cells and seeded the peripheral B_Mem_ cell compartment throughout 29 weeks. Clonal lineages with other interesting features were observed from the long GCs, including lineage 21275 which was almost exclusively IgM from week 3 to 29 (Extended Data Fig. 10b). Lastly, there were clonal lineages with clones detected in both the left ILN and right ILN at different time points during the long GCs (Extended Data Figs. 9c, 10). Out of 169 clonal lineages passing stringent criteria (HC complementarity determining region 3 [H-CDR3] length ≥14, multiple N-additions in the H-CDR3, ≥20 cells total), 11 lineages were observed in both LNs (≥10 cells per LN) at different time points, providing evidence that B_Mem_ cells generated from GCs in one LN can exit, recirculate, and enter ongoing GCs in another LN. In sum, key features observed in numerous antigen-specific clonal lineages provide direct evidence of B_GC_ cell persistence for 29 weeks, continuous accumulation of somatic mutations at substantial rates, affinity maturation, and seeding the peripheral B_Mem_ cell compartment, which together show that under select conditions, GCs are able to undergo clonal competition evolution for extremely long durations without new antigen exposure.

## Discussion

Long-lasting GCs have classically been observed in the context of chronic infections and gut microbiota exposure^24,25^, conditions known to have continuous live sources of renewed antigen. Recent reports of longer lasting GCs in influenza infection^26^, SARS-CoV-2 infection^27^, and human RNA vaccines^28–30^ have raised interest in the possibility of long-lasting GCs potentially under conditions of low or absent renewed antigen exposure. Here we demonstrate clearly that GCs can last for at least 191 days in the absence of new antigen, using an experimental system taking advantage of protein immunization, a 12-day ED immunization strategy, a robust adjuvant, and the use of Env probes to identify antigen-specific B_GC_ cells. Furthermore, we demonstrate the GCs are remarkably robust and functional for six months. The B_GC_ cells maintain proliferation, SHM, and affinity maturation, and long-lasting GCs can produce high autologous tier-2 neutralizing antibody titers, heterologous neutralizing antibody titers, and highly somatically mutated circulating Env-specific B_Mem_ cells to non-Env base epitopes.

A 12- to 14-day slow delivery (ED or osmotic pump) immunization regimen can result in substantially greater capture of vaccine antigen by stromal follicular dendritic cells (FDCs)^12^. Observation of GCs for over six months indicates that endocytic recycling of immune complexes by FDCs^31^ can be surprisingly efficient at maintaining proteins in GCs and protecting them from damage. One likely mechanism of slow delivery enhancement of GCs is improved immune complex formation, due to the supply of antigen during the earliest phases of the antibody response. Given the immunodominance of antibody responses to the non-neutralizing base of Env trimer after conventional immunizations, and epitope diversification to non-base epitopes in slow delivery immunizations (Fig. 2g), we speculate that immune complexes with Env under slow delivery conditions are primarily composed of base-binding antibodies, which shield the base of the Env trimer in GCs and orient the trimers to better display neutralizing epitopes antipodal to the base, thereby enriching for neutralizing antibody B cells. This is further illustrated by the shift away from base directed immunodominance in B_Mem_ cells during the long prime (Fig 4i), as opposed to a normal 10-week boost that appears to recall more base-specific B cells. Thus, the improved autologous and heterologous neutralization by Group 3 animals is likely partly owing to the diversity of B cells recruited and partly due to increased affinity maturation from extensive GC responses.

We have shown that GCs can persist for greater than six months in response to a priming immunization, with a number of notable outcomes. These findings indicate that patience can have great value for allowing antibody diversification and evolution in GCs over surprisingly extended periods of time. A long prime, adjuvanted, escalating dose immunization approach holds promise for difficult vaccine targets.

## METHODS

### Protein expression and purification

BG505 MD39 SOSIP Env trimers (MD39) were co-expressed with furin in HEK293F cells and expressed as previously described^14^. Trimers used for immunizations were expressed tag-free and quality checked for low endotoxin levels. BG505 MD39 SOSIP and BG505 MD39-base knockout (KO) trimers used as baits in flow cytometry were expressed with a C-term Avi-tag and biotinylated using a BirA biotinylation kit according to manufacturer’s instructions (Avidity). The BG505 MD39-base KO trimer had the following mutations relative to the BG505 MD39 SOSIP: A73C, R500A, P561C, C605T, S613T, Q658T, L660N, A662T, and L663C.

### Animals and immunizations

For MD39 plus SMNP ED immunization groups, Indian rhesus macaques (RMs, *Macaca mulatta*) were housed at Alpha Genesis Inc. and treated in accordance with protocols approved by the Alpha Genesis Inc. Animal Care and Use Committee (IACUC). 2 females and 2 males with matched aged and weight were assigned to each experimental group. Animals were 2-3 years old at the time of the priming immunization. All immunizations were given subcutaneously (s.c.) in the left and right mid-thighs with a total dose of 50 µg MD39 and 375 µg SMNP each side. For priming, a 12-day escalating dose strategy was used (Extended Data Fig. 1a)^12^.

For the MD39 plus alum bolus group, RMs were housed at the Tulane National Primate Research Center as part of a larger NHP study (Phung et al. Manuscript in preparation). This study was approved by the Tulane University IACUC. Animals were grouped together to match age, weight, and gender. Animals were between 3.5–5 years at time of first immunization, with 3 females and 3 males in the study group. All immunizations were given s.c. in the left and right mid-thighs with 50 µg MD39 and 500 µg alum (Alhydrogel adjuvant 2%; InvivoGen) per side. All animals were maintained in accordance with NIH guidelines.

### Lymph node fine needle aspiration

LN FNAs were used to sample the left and right inguinal LNs and performed by a veterinarian. Draining lymph nodes were identified by palpation. A 22-gauge needle attached to a 3 cc syringe was passed into the LN up to 5 times. Samples were placed into RPMI containing 10% fetal bovine serum, 1X penicillin/streptomycin (pen/strep). Samples were centrifuged and Ammonium-Chloride-Potassium (ACK) lysing buffer was used if the sample was contaminated with red blood cells. Samples were frozen down and kept in liquid nitrogen until analysis.

### Flow cytometry and sorting

Frozen FNA or PBMC samples were thawed and recovered in 50 % (v/v) FBS in RPMI. Recovered live cells were enumerated and stained with the appropriate staining panel. MD39 and MD39-base KO baits were prepared by mixing biotinylated MD39 with fluorophore-conjugated streptavidin (SA) in small increments at RT in an appropriate volume of 1X PBS over the course of 45 min. MD39:SA were added to the cells for 20 minutes, after which the antibody master mix was added for another 30 minutes at 4 ºC. Where KO baits were used, KO baits were added to the cells first for 20 minutes, then WT MD39:SA baits and were added for another 20 min, followed by the addition of the remainder of the staining panel for an additional 30 minutes at 4ºC, similar to previously described^12^. Fully supplemented RPMI (R10; 10 % (v/v) FBS, 1x pen/strep, 1x Glutamax) was used as FACS buffer. For sorting, anti-human hashtag antibodies (Biolegend) were individually added to each sample at a concentration of 2.5 µg/ 5 million cells at the time of addition of the master mix. Group 1 samples were sorted on a FACSFusion (BD Biosciences) and Group 2 and 3 samples were either acquired or sorted on a FACSymphony S6 (BD Biosciences). Indexed V(D)J, Feature Barcode, and GEX libraries of sorted samples were prepared according to the protocol for Single Indexed 10X Genomics V(D)J with Feature barcoding kit (10X Genomics). Custom primers were designed to target RM BCR constant regions. Primer set for PCR 1; forward: AATGATACGGCGACCACCGAGATCTACACTCTTTCCCTACACGACGCTC, reverse: AGGGCACAGCCACATCCT, TTGGTGTTGCTGGGCTT, TGACGTCCTTGGAAGCCA, TGTGGGACTTCCACTGGT, TGACTTCGCAGGCATAGA. Primer set for PCR 2; forward: AATGATACGGCGACCACCGAGATCT, reverse: TCACGTTGAGTGGCTCCT, AGCCCTGAGGACTGTAGGA, AACGGCCACTTCGTTTGT, ATCTGCCTTCCAGGCCA, ACCTTCCACTTTACGCT. Forward primers were used at a final concentration of 1 µM and reverse primers at 0.5 µM each per 100 uL PCR reaction. Libraries were pooled and sequenced on a NovaSeq Sequencer (Illumina) as previously described^32^.

During the long prime tracking phase (weeks 16-25, right LNs), samples were stained as described above, fixed in BD Cytofix (BD Biosciences), then analyzed on a FACSCelesta (BD Biosciences). For intracellular staining, cells were stained as described above then fixed with Foxp3 / Transcription factor staining kit (Invitrogen). Cells were washed with 1x diluted permeabilization buffer then stained for 1 hr with antibodies targeting transcription factors of interest. Cells were washed and analyzed on a Cytek Aurora (Cytek Biosciences). All flow cytometry data were analyzed in Flowjo v10 (BD Biosciences).

For LN FNA data inclusion for GC gating for MD39 plus alum (Group 1), a threshold of 250 total B cells in the sample was used. For Env-binding GC B cell gating, a threshold of 75 total GC B cells was used. Any sample with fewer than 75 GC B cells but with a B cell count of > 500 cells was set to a baseline of 0.001% Env^+^ GC B cells (% of B). Otherwise the limit of detection was calculated based on the median of [3/(number of B cells collected)] from the pre-immunization LN FNA samples.

The following reagents were used for staining: Alexa Fluor 647 SA (Invitrogen), BV421 SA (Biolegend), BV711 SA (Biolegend), PE SA (Invitrogen), Live/Dead fixable aqua (Invitrogen), Propidium iodide (Invitrogen), eBioscience Fixable Viability Dye eFluor 780 (Invitrogen), mouse anti-human CD20 BV785, BUV395, Alexa Fluor 488, PerCP-Cy5.5 (2H7, Biolegend), mouse anti-human IgM PerCP-Cy5.5, BV605 (G20-127, BD Biosciences), mouse anti-human CD4 BV650, Alexa Fluor 700 (OKT4, Biolegend), mouse anti-human PD1 BV605 (EH12.2H7, Biolegend), mouse anti-human CD3 BV786, APC-Cy7 (Sp34-2, BD Biosciences), mouse anti-human CXCR5 PE-Cy7 (MU5UBEE, ThermoFisher), mouse anti-human CD71 PE-CF594 and FITC (L01.1), mouse anti-human CD38 PE, APC (OKT10, NHP Reagents), mouse anti-human CD8a APC eFluor 780 (RPA-T8, ThermoFisher), mouse anti-human CD14 APC-Cy7 (M5E2, Biolegend), mouse anti-human CD16 APC-Cy7 (3G8, Biolegend), mouse anti-human CD16 APC-eFluor 780 (ebioCD16, Invitrogen), mouse anti-human IgG Alexa Fluor 700, BV510, and BV786 (G18-145, BD Biosciences), Mouse anti-NHP CD45 BUV395 (D058-1283, BD Biosciences), mouse anti-human BCL6 Alexa Fluor 647 (K112-91, BD Biosciences), mouse anti-human KI67 BV480 (B56, BD Biosciences), mouse anti-human FoxP3 BB700 (236A/E7, BD Biosciences), mouse anti-human CD27 PE-Cy7 (O323, Biolegend), goat anti-human IgD FITC (polyclonal, Southern Biotech), Armenian hamster anti-mouse/human Helios PE/Dazzle 594 (22F6, Biolegend), TotalSeq-C anti-human Hashtag antibody 1-8 (LNH-94 and 2M2, Biolegend), TotalSeq-C0953 PE Streptavidin (Biolegend).

### Detection of antigen-specific GC-T_FH_ cells

Antigen induced marker (AIM)-based identification of Env-specific GC-T_FH_ cells was performed as previously described.^12,16^ In summary, cells were thawed in 50 % (v/v) FBS in RPMI and resuspended in 500 µL of DNase in R10 (100 µL DNAse in 900 µL R10) for 15 min at 37ºC in a CO_2_ and humidity controlled incubator. 5 mL R10 was added and cells were further rested for 3 hrs. Cells were enumerated and seeded at ∼1 million cells per well in R10, and incubated with a final concentration of 2.5 µg/mL MD39 Env peptide pool, 10 pg/mL SEB, or media only (unstimulated) for 18 hrs at 37ºC in a CO2 and humidity controlled incubator. 1:100 mouse anti-human CXCR5 PE-Cy7 (MU5UBEE, ThermoFisher) was added to each well at the start of stimulation. Cells were washed and stained for 45 min in the dark at 4 ºC. After staining, cells were washed and fixed with BD Cytofix (BD Biosciences) and analyzed on a BD FACSCelesta (BD Biosciences). The following antibodies were used in the flow panel: mouse anti-human CD4 Alexa Fluor 700 (OKT4, Biolegend), mouse anti-human CD20 BV785 (2H7, Biolegend), mouse anti-human PD1 BV605 (EH12.2H7, Biolegend), mouse anti-human CXCR5 PE-Cy7 (MU5UBEE, ThermoFisher), mouse anti-human CD134 PE (L106, BD Biolegend), mouse anti-human 4-1BB APC (4B4-1, Biolegend), mouse anti-human CD25 FITC (BC96, Biolegend), mouse anti-human CD16 APC-eFluor 780 (ebioCD16, Invitrogen), mouse anti-human CD8a APC eFluor 780 (RPA-T8, ThermoFisher), mouse anti-human CD14 APC-Cy7 (M5E2, Biolegend), eBioscience Fixable Viability Dye eFluor 780 (Invitrogen).

### Neutralization assays

Pseudovirus neutralization assays were performed as previously described^11^. BG505 pseudovirus neutralization was tested using the BG505.W6M.ENV.C2 isolate with the T332N mutation to restore the N332 glycosylation site, except in Extended Data Fig. 3a (Duke) and 3b, where the original T332 strain was used. Heterologous neutralization breadth was tested on a panel of 12 cross-clade isolates, representative of larger virus panels isolated from diverse geography and clades^33^. The cut-off for neutralizing serum dilution was set at 1:30 or 1:20 depending on the starting serum dilution. Absolute ID_50_s were calculated using normalized RLU values and a customized non-linear regression model:

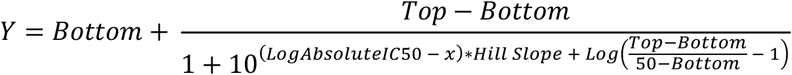

with the bottom constraint (Bottom) set to 0 and top constraint (Top) set to <100 model in Prism 8 (GraphPad).

### ELISA

Plasma samples were thawed, heat inactivated at 56ºC for at least 30 min, and spun down. 96-well half area plates (Corning) were coated overnight with SA at 2.5 µg/mL. Plates were washed 3x with wash buffer (PBS, 0.05% (v/v) Tween-20), then coated with biotinylated MD39 or MD39-base KO trimmers at 1 µg/mL. After washing 3x with wash buffer, plates were blocked with blocking buffer (PBS, 3% (w/v) BSA) for 1hr at RT. Plasma serially diluted in blocking buffer were allowed to bind the trimers for 1 hr at RT. Plates were washed 3x and incubated with goat anti-rhesus IgG-HRP antibody (Southern Biotech, 1:10,000 in blocking buffer) for 1 hr at RT. Plates were washed 6x, and developed with 1-Step Ultra TMB (ThermoFisher). The reaction was stopped with an equivalent volume of 2N H_2_SO_4_ (Ricca Chemical Company) and signal was read at OD 450 nm on an EnVision plate reader (Perkin Elmer). Endpoint titers were interpolated from a Asymmetric Sigmoidal, 5PL X is log(concentration) model in Prism 9 (GraphPad).

### EMPEM Analysis

Polyclonal EM analysis was performed as previously described^19,34^. Plasma antibodies were purified using Protein A Sepharose resin (GE Healthcare), eluted from the resin with 0.1 M glycine at pH 2.5 and buffer exchanged into 1X PBS. Fabs were generated using crystalline papain (Thermo Scientific) and digested for 5 h at 37 °C, and purified via size exclusion chromatography (SEC) using Superdex 200 Increase 10/300 column (GE Healthcare). Complexes were assembled with 0.5 mg of polyclonal Fabs incubated overnight with 15 µg of MD39 Env trimers at RT, followed by purification to remove unbound Fab via size exclusion chromatography (SEC) using a Superose 6 Increase 10/300 column (GE Healthcare). Complexes were diluted to 30-50 µg/mL and immediately placed on 400-mesh Cu grids and stained with 2% (w/v) uranyl formate for 40 s. Images were collected via the Leginon automated imaging interface using either a Tecnai Spirit electron microscope, operating at 120 kV, or a Tecnai TF20 electron microscope operated at 200 kV. For the Spirit, nominal magnification was 52,000x, with a pixel size of 2.06 Å. The TF20 was operated at a nominal magnification of 62,000x with a pixel size of 1.77 Å for the TF20. Micrographs were recorded using a Tietz 4k × 4k TemCam-F416 CMOS camera. Particles were extracted via the Appion data processing package^35^ where approximately 100,000 particles were auto-picked and extracted. Using Relion 3.0^36^, particles were 2D-classified into 100 classes and particles with antigen-Fab characteristics were selected for 3D analysis. Initially, 3D classification was done using 20-40 classes, with a low-resolution model of a non-liganded HIV Env ectodomain used as a reference. Particles from similar looking classes were combined and reclassified, with a subgroup of 3D classes processed using 3D auto-refinement. UCSF Chimera 1.13 was used to visualize and segment the 3D refined maps.

### BCR sequencing and processing

A custom RM germline VDJ library was generated using references published by Cirelli et al.^12^, and Bernat et al^37^. CellRanger V3.0 was used to assemble full length V(D)J reads. The constants.py file in the CellRanger VDJ python library was modified to increase the maximum acceptable CDR3 length to 110 NT. CellRanger V6 was used to obtain gene expression counts from sequenced GEX libraries. Libraries were aligned to the Ensemble Mmul10 reference genome, with addition of mitochondrial genes from Mmul9. Sequences were demultiplexed by hashtags using the MULTIseqDemux command in Seurat V4.^38^ For HC sequences where both kappa and lambda LC contigs were detected, the B cell was assigned a lambda LC, because lambda LC rearrangement only takes place if the kappa LC is not productive.

### Longitudinal lineage and somatic mutation analysis of BCR sequences

VDJ sequence output from CellRanger was further analyzed using packages from the Immcantation portal^39^. An IgBLAST database was built from the custom RM germline VDJ Library. This was then used to parse the 10X V(D)J output from CellRanger into an AIRR community standardized format using the Change-O pipeline to allow for further downstream analysis with the Immcantation portal. Clonal lineages were determined for each animal with DefineClones.py, using the appropriate clustering threshold as determined by the distToNearest command from the SHazaM package in R. Inferred germline V and J sequences from the reference library were added with CreateGermline.py. Because germline D genes sequences and N nucleotide additions cannot be accurately predicted, these were masked from further analysis. The total number of mutations (V and J-genes) for each HC and LC sequence was calculated by counting the number of nucleotide changes between the observed and predicted germline sequences with SHazaM’s observe mutation command. For the analysis of total HC mutations, all productive HC contigs were analyzed. For LCs, only contigs paired with HCs were assessed. Sequences where the VH or VL call aligned to alleles IGHV3-100*01, IGHV3-100*01_S4205, IGHV3-100*01_S4375, IGHV3-36*01_S5206, IGHV3-36*01_S6650, IGHV3-NL_11*01_S5714, IGHV4-79-a, IGHV4-NL_1*01_S0419, IGLV1-69, IGLV1-ACR*0 or IGLV2-ABX*01, were found to have an extremely high degree of substituted nucleotides at all timepoints compared to their inferred germline sequences, likely because of poor V gene assignment due to an incomplete V(D)J reference library. These sequences were excluded from further analysis. Only paired HC-LC BCR sequences were analyzed when building clonal trees. Maximum-likelihood lineage trees were built for clonal families with Dowser^40^ using the pml method in the GetTrees function. For lineage trees, the length of the branches represents the estimated number of total mutations that have occurred in each HC clone and its most recent common ancestor in lineage, rather than a simple count of nucleotide changes in the germline sequence.

### Transcriptomics analysis

The package Seurat V4^38^ was used for graph-based clustering and visualizations of the gene expression data generated by CellRanger. Initial filtering was conducted on each sample to remove cells expressing < 200 or > 4500 genes as well as cells with >10% of their transcriptome made up of mitochondrial DNA. Gene expression counts were log normalized via the NormalizeData command. A list of common variable genes for across all sample were identified with the SelectIntergrationFeatures function. Expression of these common variable genes was scaled using Principal component analysis (PCA) conducted with with RunPCA. Next all samples were integrated together into a single dataset using an RPCA reduction to remove batch effects via the FindIntegrationAnchors and IntergrateData commands. Louvian clustering was conducted on the entire integrated dataset with the FindNeighbours and FindCluster functions. Clusters containing large numbers of cells with high levels of mitochondrial DNA as well as clusters with low *MS4A1* (CD20) and *CD19* expression were excluded from further analysis. Differentially expressed genes were identified in Seurat with the FindMarkers function by running Wilcoxon rank sum test for each cluster against all other clusters. Gene Set Enrichment Analysis (GSEA) was conducted using the fgsea package in R^41,42^. Differentially expressed genes from previously identified human LZ, DZ, Intermediate, PreMem and Plasmablast subsets were taken from Holmes et al.^23^ and combined to create the gene sets used for GSEA.

### Graphs, Statistics, and Cell Generation Calculation

All statistics were calculated in Prism 9 or R. The statistical tests used are indicated in the respective figure legends and utilize a two-tailed test. All graphs were generated in Prism 9 or R. Geometric mean and geometric standard deviation (SD) are shown for data plotted on a Log_10_ axis. Mean and SD are plotted for data graphed on a linear axis. Median and quartiles 1 and 3 are shown for violin plots. UMAP plots were generated using Seurat V4^38^. For comparison of total HC and LC mutations between Group 2 and Group 3, statistical significance was calculated only if mean mutations were significantly different by per animal comparisons.

Chao1 estimation of clonal richness was calculated according to the following formula:

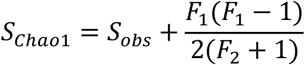

Where S_obs_ is the total number of observed species and F_1_ and F_2_ are the number of singletons and doubletons, respectively.

## ACKNOWLEDGEMENTS

We thank the Flow Cytometry Core at the La Jolla Institute for Immunology (LJI) and LJI Sequencing Core. We also thank Gunilla Karlsson Hedestam and Martin Corcoran from the Karolinska Institutet for providing additional naive RM IGHV reference sequences prior to publication. This study was partially performed at AlphaGenesis Inc. Funding: This work was supported in part by the National Institute of Allergy and Infectious Diseases of the NIH under award numbers CHAVD-ID UM1AI100663, CHAVD UM1AI144462, R01 AI125068, R01 AI136621, P01AI048240, and NIH NIAID SVEU Contract No. HHSN272201300004I. This work was also supported by the Bill and Melinda Gates Foundation Collaboration of AIDS Vaccine Discovery (CAVD) OPP1115782/INV-002916.

## DATA AVAILABILITY

Data will be deposited upon acceptance of the manuscript. RNA-seq data will be deposited in the Gene Expression Omnibus database. BCR sequences have will be deposited in Genbank. 3D EM reconstructions will be deposited into the Electron Microscopy Data Bank. Sequencing data and EM particle stacks are available upon request.

## AUTHOR INFORMATION

### Contributions

J.H.L and S.C conceived and designed experiments. D.J.I provided conceptual insights. J.H.L performed and analyzed most experiments pertaining to LN FNA data, generated FNA BCR sequencing data, and assisted in FNA BCR sequencing analysis. H.S. analyzed BCR data, performed all experiments related to PBMC samples, generated and analyzed single cell transcriptomics data. J.H.L and C.K performed serum ELISAs. R.N and L.H performed neutralization assays and analysis; S.S and D.M performed independent confirmation neutralization assays and analysis. S.R, L.S, J.T, W.L, and G.K.O generated and analyzed EMPEM data. D.J.I and M.S supplied the SMNP adjuvant. C.A.C, E.G, M.K, S.H, T.-M.M, Y.A, and W.R.S supplied immunogens and flow cytometry baits. C.C and K.C designed primers. I.P, A.K, C.A-H, M.F, B.G, J.D, F.S, P.P.A, and R.S.V performed MD39 plus alum rhesus studies. D.G.C and G.S provided technical guidance for studies performed at the Tulane National Primate Research Center. A.B.W, and D.R.B, provided supervision. J.H.L and S.C. wrote the original draft. H.S and L.H contributed to figure generation. H.S, D.R.B, D.J.I, A.B.W, W.R.S provided comments and assisted in revision.

