## Extended Figures 1-10 for "Long-lasting germinal center responses to a priming immunization with continuous proliferation and somatic mutation"

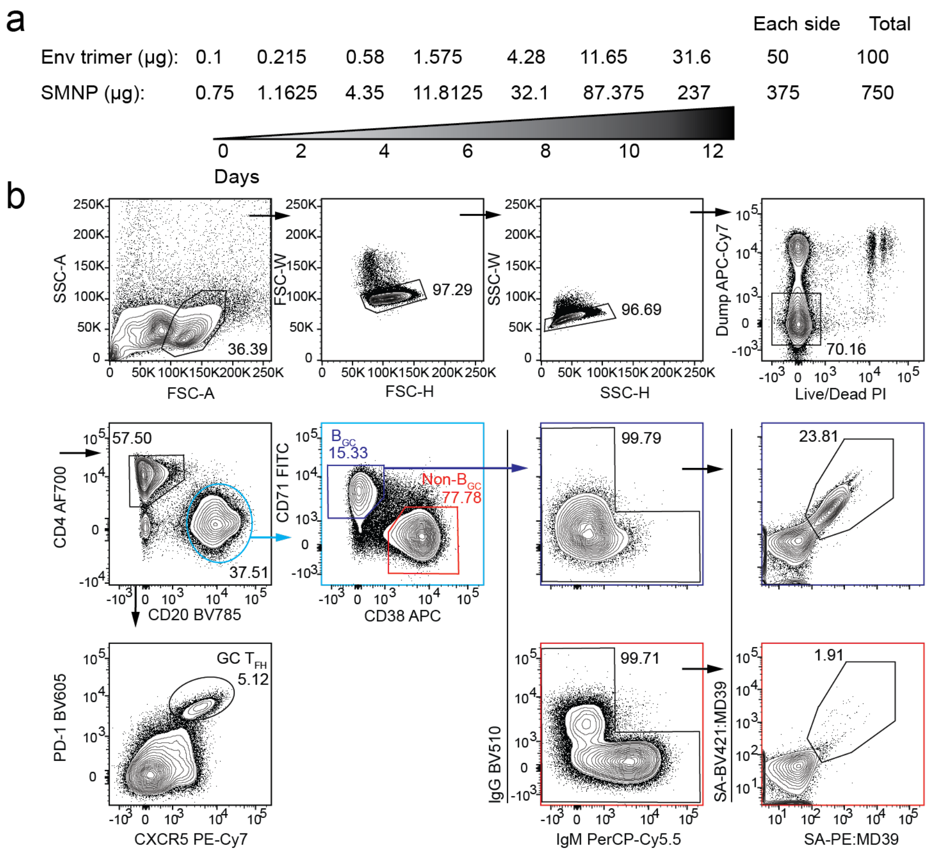

**Extended Data Fig 1: Escalating dose immunization strategy and representative flow cytometry analysis of FNA samples. a,** Priming ED strategy and injection schedule. **b,** Gating strategy for the longitudinal analysis of GC-$T_{FH}$ and $B_{GC}$ cells from ILN FNA samples. $CD71^{+}CD38^{-}$/ $MD39^{+/+}$ cells were sorted for BCR sequencing and transcriptomic analyses. $CD71^{+}CD38^{-}/MD39^{-/-}$ cells were also sorted for weeks 3,4,7, and 10.

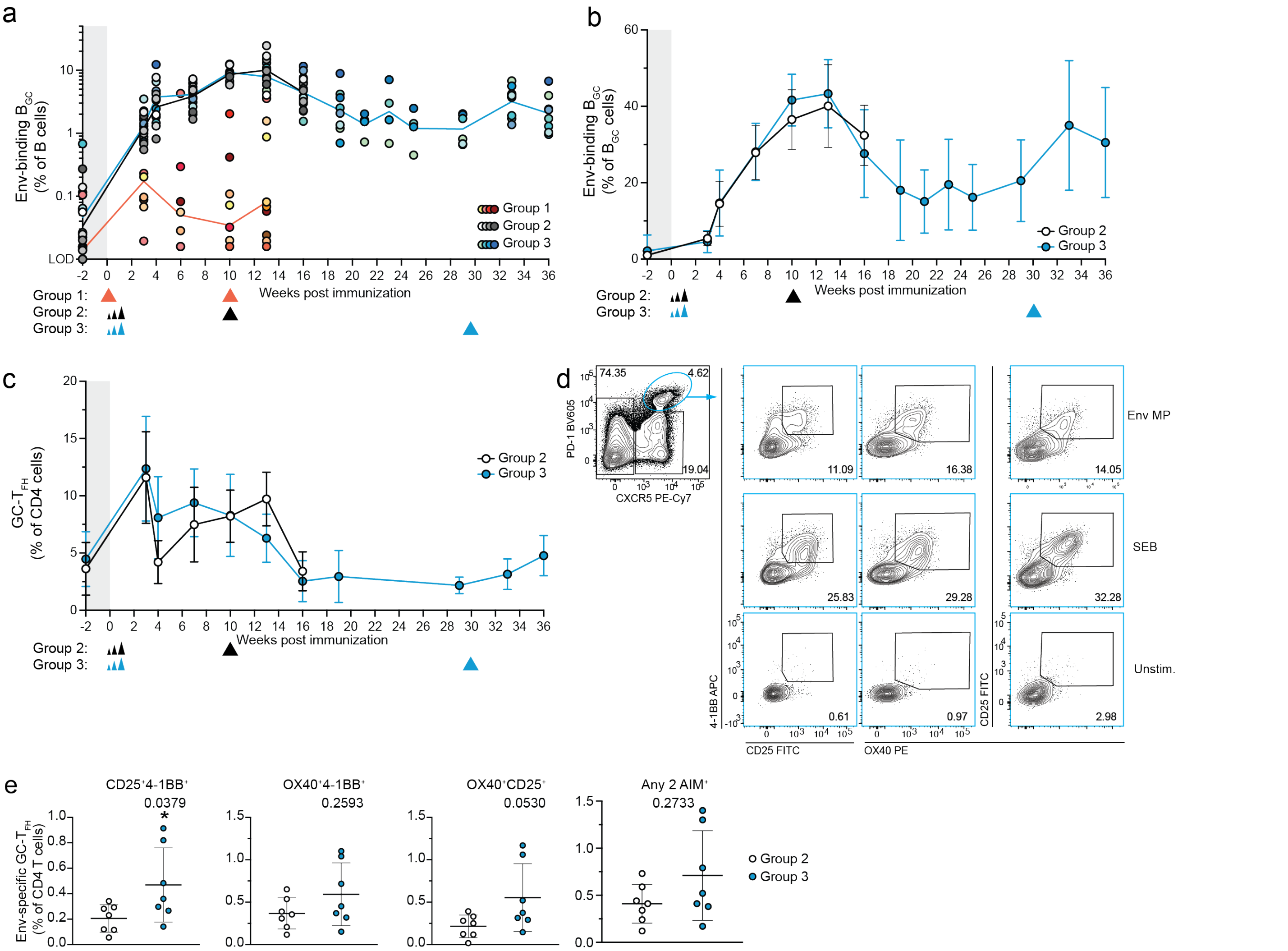


**Extended Data Fig 2: $B_{GC}$ and $GC\text{-}T_{FH}$ kinetics. a,** Env-binding $B_{GC}$ cells as a percent of total $CD20^{+}$ B cells. **b,** Env-binding $B_{GC}$ cells as a percent of total $B_{GC}$ cells. **c,** Kinetics of $GC\text{-}T_{FH}$ cells for ED/SMNP immunized animals. Gating strategy is shown in Extended Data fig. 1d. **d,** Representative flow plot of an AIM assay to detect Env-specific $GC\text{-}T_{FH}$ cells after 18 hrs of ex vivo restimulation. Stimulation conditions are indicated on the right: 15-mer overlapping Env peptide megapool (Env MP), staphylococcal enterotoxin (SEB), or unstimulated (unstim.). **e,** Env-specific $GC\text{-}T_{FH}$ cells ($CXCR5^{hi}PD\text{-}1^{hi}$, gated on FSC-A/SSC-A, FSC-H/FSC-W, SSC-H/SSC-W, Live/Dead Fixable $e780^{-}$, CD4 $AF700^{+}$B220 $BV785^{-}$) detected at 6 wks post boost (Group 2: week 16, Group 3: week 36) quantified by AIM assay using the indicated pair of activation markers, or positive for any two of the three AIM markers (Or gates). Data shown is background ($AIM^{+}$ signal from unstimulated samples) subtracted. Left and right ILNs are graphed as independent data points. Mean and SD or geometric mean and geometric SD are plotted depending on the scale. Limit of detection (LOD). Mann-Whitney test: ns > 0.05; *$p < 0.05$.

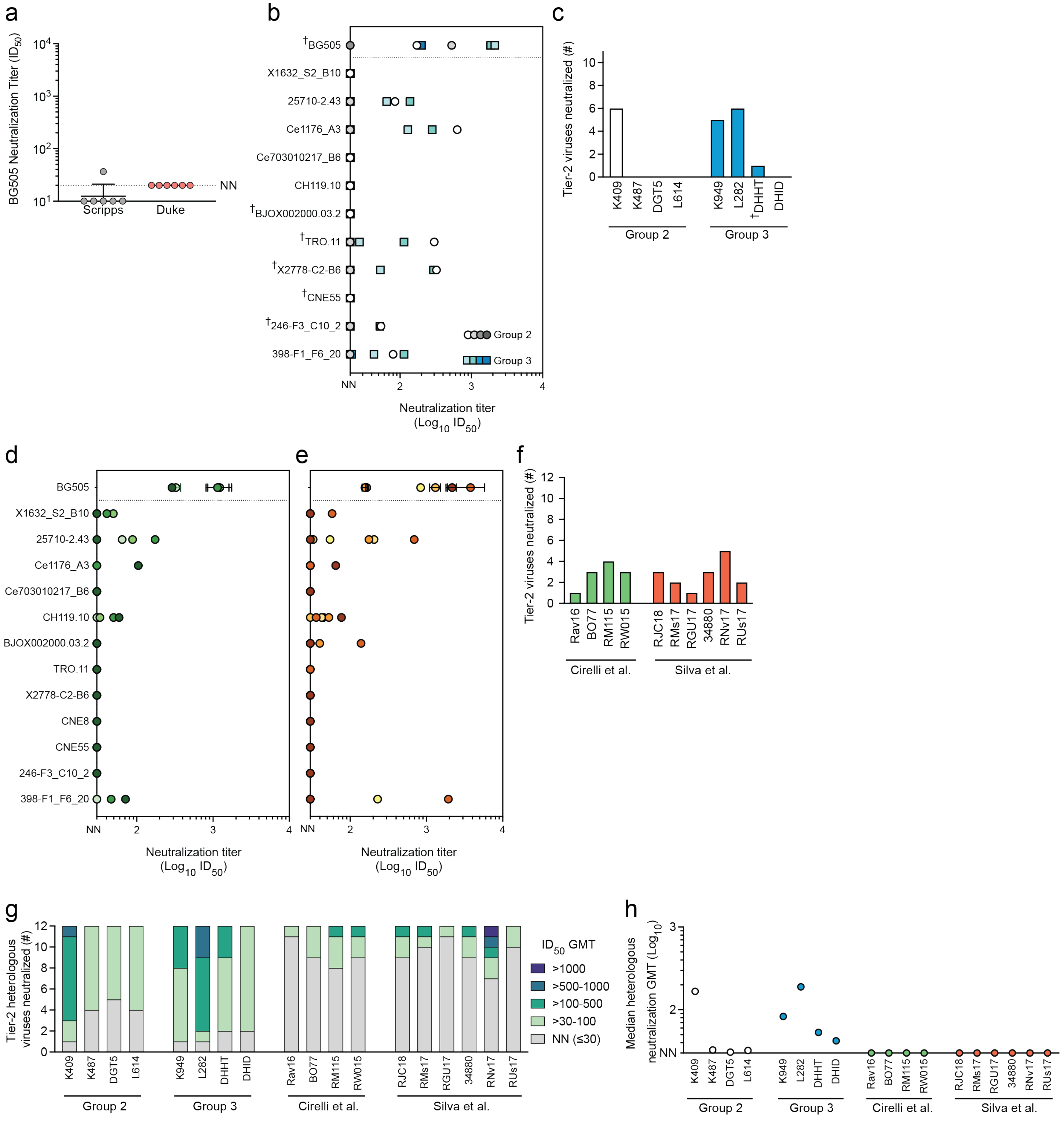


**Extended Data Fig 3. DE immunization using the SMNP adjuvant improves the quality of Env-specific antibody responses. a,** Group 1 autologous neutralization titers determined by assays performed by two independent labs (Scripps Research Institute [Scripps] or Duke). $ID_{50}$ < 20 is considered non-neutralizing (NN). **b,** Week 3 post-boost Group 2 and 3 serum neutralization $ID_{50}$ titers determined independently from a second laboratory (Duke). $ID_{50}$ cut-off for NN < 20. †Viruses for which serum neutralization was not determined for Group 3 animal DHHT. **c,** Number of heterologous tier-2 viruses neutralized out of 11 tested in (b). †Neutralization for only 6 viruses was tested for DHHT. **d-e,** Serum neutralization assays (Scripps) tested head-to-head with data shown in Fig. 2e using week 3 post-boost 2 sera from four RMs ED immunized with an ISCOM adjuvant (SMNP without MPLA) + BG505 (d)[12], and week 2 post-boost 2 sera from six RMs bolus immunized with SMNP + MD39 (e)[15]. $ID_{50}$ cut-off for NN < 30. **f,** Number of tier-2 heterologous viruses neutralized by post-boost 2 serum tested in d-e[12,15]. **g,** Number of tier-2 heterologous viruses neutralized with the indicated serum GMT titers (Scripps). **h,** Median tier-2 heterologous neutralization GMT across the 12-virus panel (Scripps).

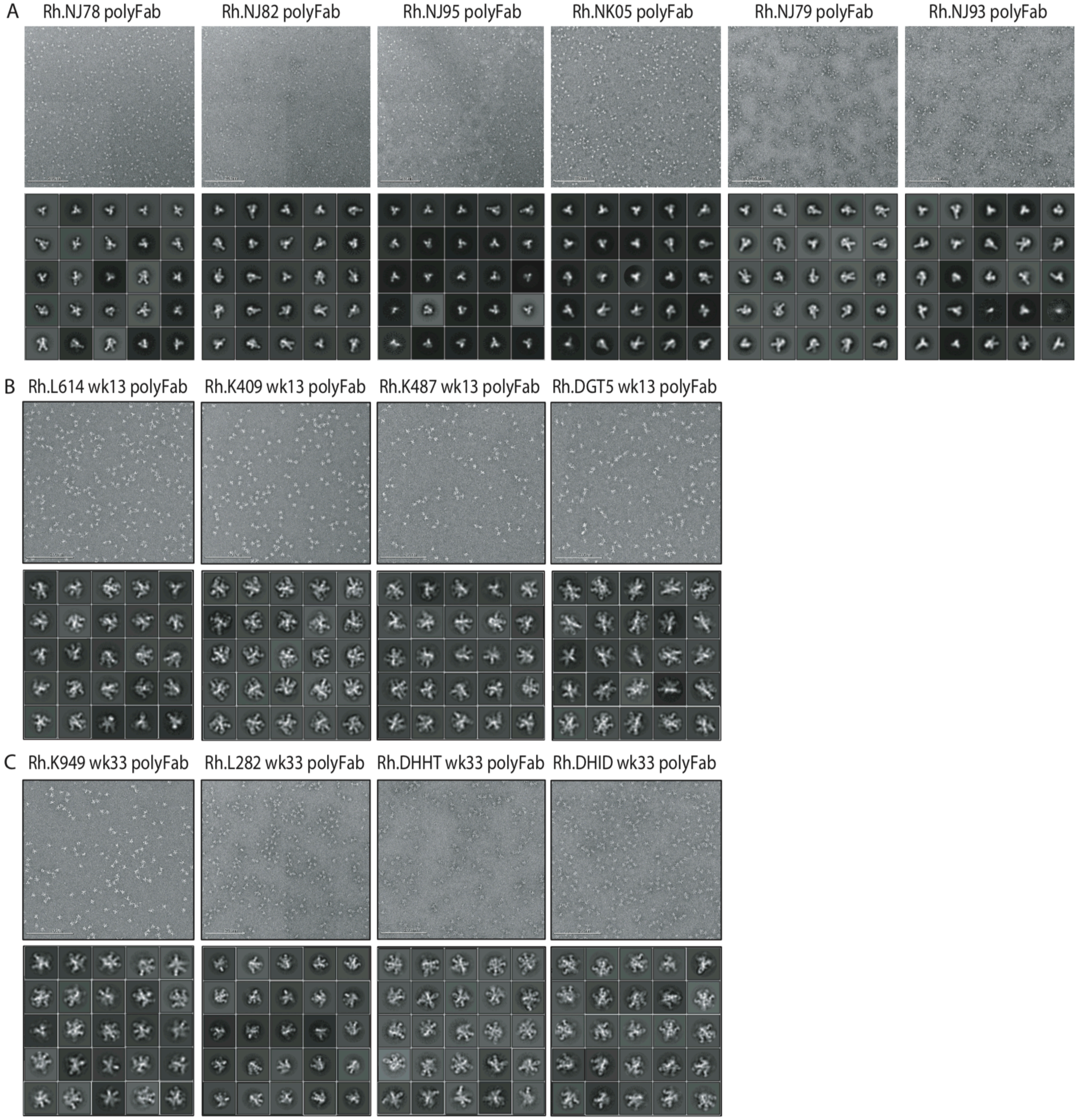


**Extended Data Fig 4. EMPEM analysis of polyclonal antibodies. a-c,** negatively stained EM micrographs of MD39 Env trimer: polyclonal Fab complexes (top), and 2D-class averages (bottom). Data for RMs in Group 1 (a), Group 2 (b), and Group 3 (c). The scale bar shown in the lower left corner of each micrograph corresponds to 200 nm.

a

| Animal ID | LN | Weeks | % Env+ $B_{GC}$ of B cells | |
|---|---|---|---|---|
| | | | CD71⁺CD38⁻ Env⁺/⁺ | KI67⁺BCL6⁺ Env⁺/⁺ |
| DHHT | L | 21 | 0.57 | 0.54 |
| DHID | L | 25 | 0.23 | 0.20 |
| K949 | L | 21, 23, 25 combined | 0.48 | 0.40 |
| L282 | L | 21 | 0.39 | 0.17 |
| L282 | L | 21, 23 combined | 0.49 | 0.32 |

b

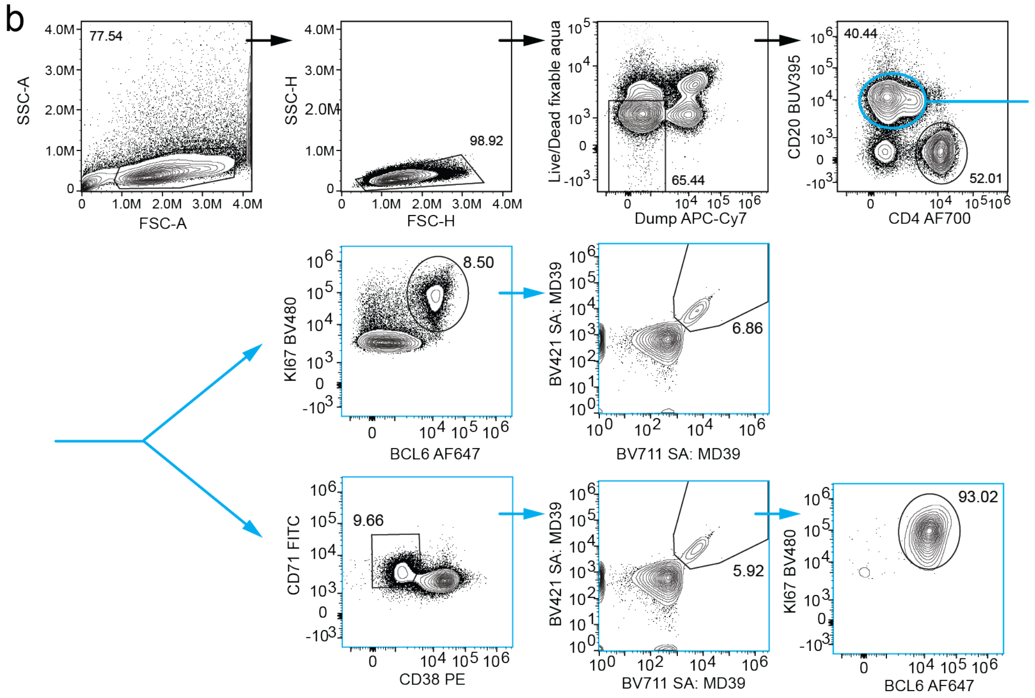


**Extended Data Fig 5. Intracellular staining of $B_{GC}$ markers. a,** Some left ILN samples were pooled for the intracellular staining panel, for higher cell numbers. **b,** Representative flow plot and complete gating strategy for BCL6 and KI67 staining.

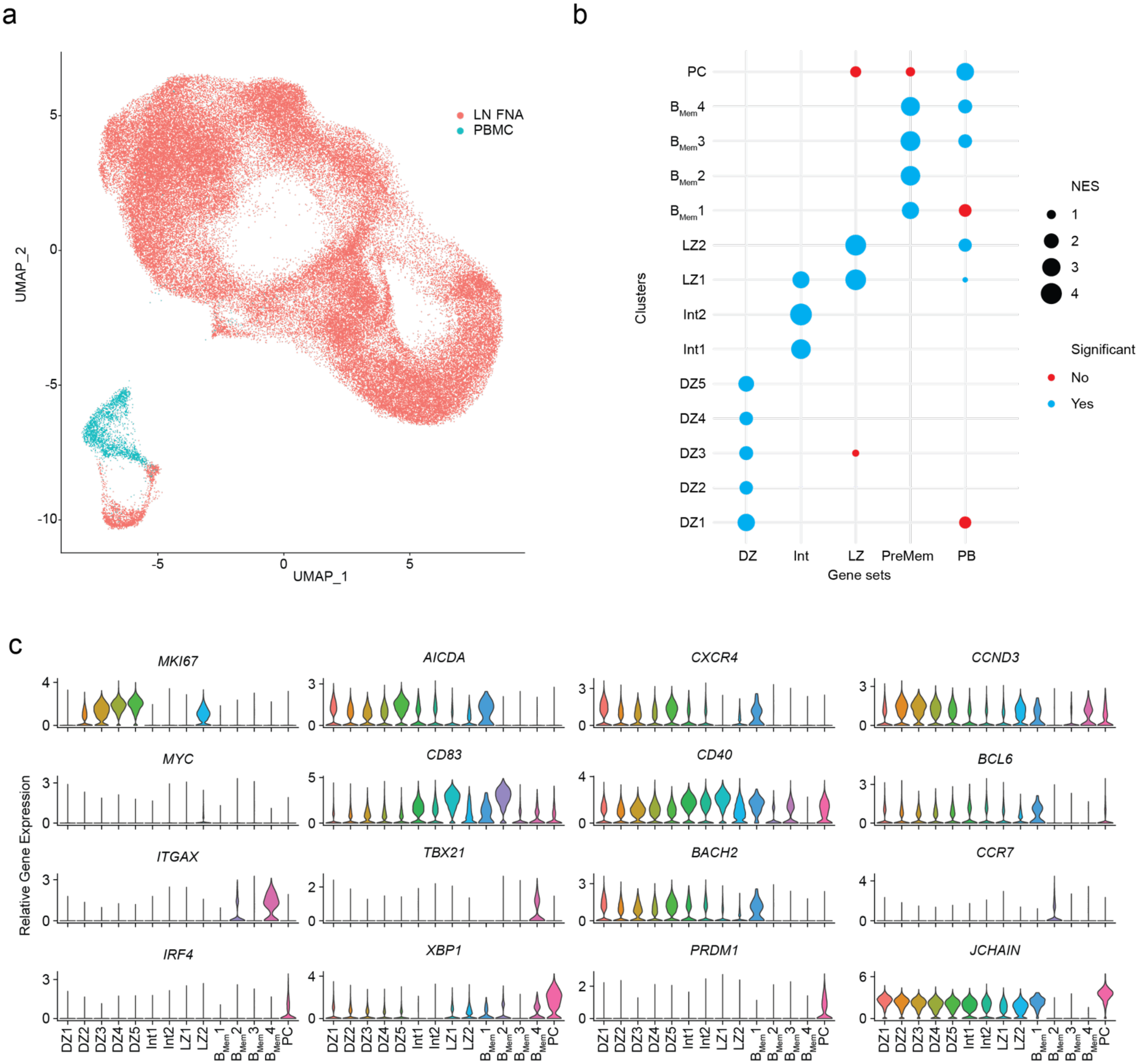


**Extended Data Fig 6. Single cell transcriptional profiling of B cells. a,** Single cell transcripts of LN FNA $B_{GC}$ (Group 2 and 3 weeks 3, 4, 7,10 and 16, Group 2 week 13, Group 3 weeks 29 and 33) and PBMC $B_{Mem}$ cells (Group 2 week 16, Group 3 week 36) were assessed. **b,** A summary of the results from GSEA of upregulated gene profiles from single cell clusters using previously identified B cell subset gene signatures[23]. Size of the dots represent the normalized enrichment score (NES). Significant NES results are shown in blue, nonsignificant results in red. **c,** Expression levels of the displayed genes used to help identify clusters of B cell subsets.

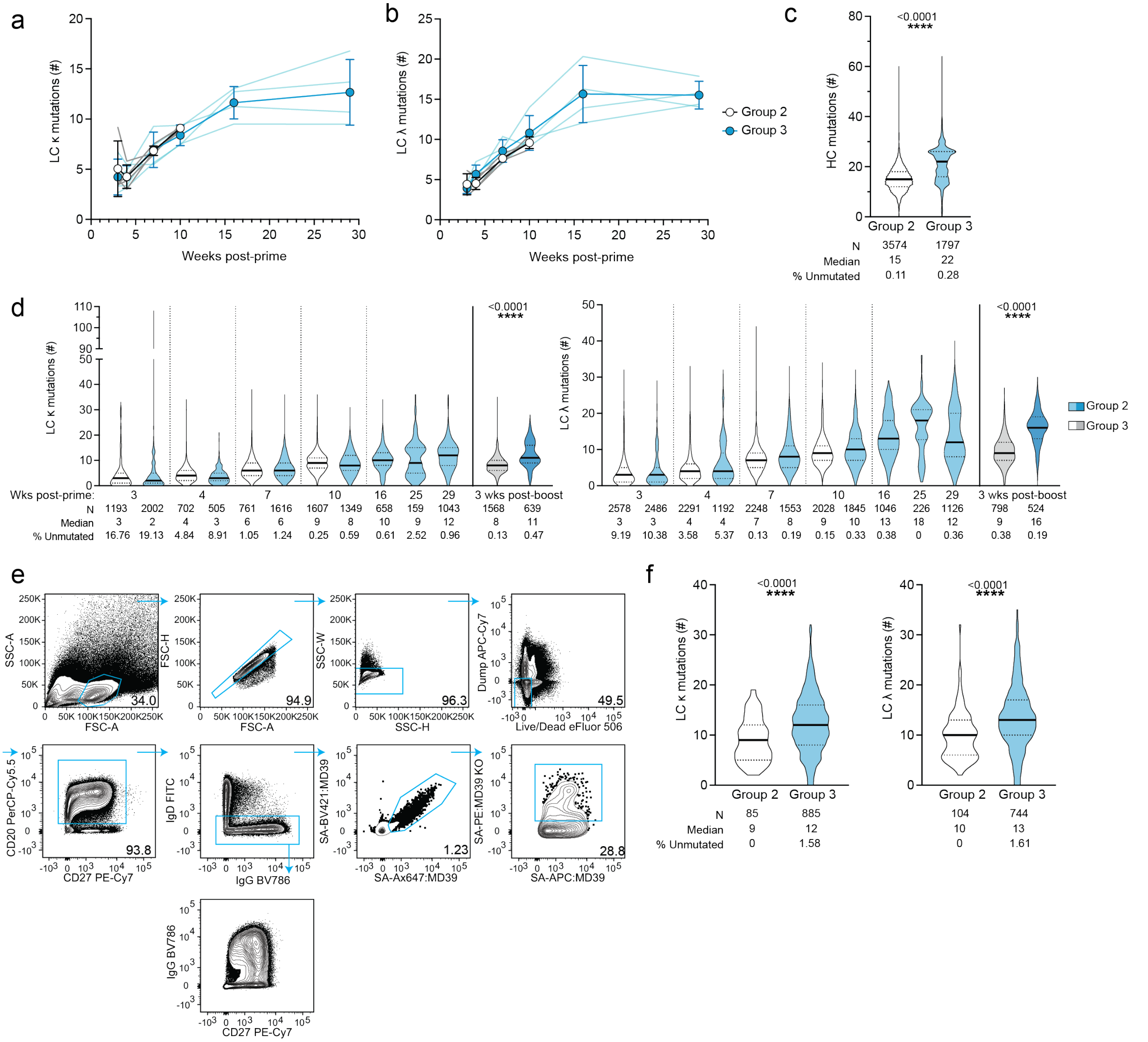


**Extended Data Fig 7. BCR sequences of $B_{GC}$ and $B_{Mem}$ cells. a-b,** Number of NT mutations in the V and J-gene region of LC sequences (Kappa: LC κ, Lambda: LC λ) derived from Env-specific $B_{GC}$ cells after priming. Spaghetti plots track the number of mutations in each animal. **c,** Number of HC mutations at week 6 post-boost. **d,** Number of LC κ and LC λ NT mutations in $B_{GC}$ cells, respectively. There are two outliers among Group 3 week 3 LC κ sequences, each with 108 and 105 mutations. **e,** Gating strategy for the detection of Env-specific $B_{Mem}$ cells. **f,** Number of LC κ and LC λ NT mutations in wk 6 post-boost $B_{Mem}$ cells. Mann-Whitney test, ****P < 0.0001.

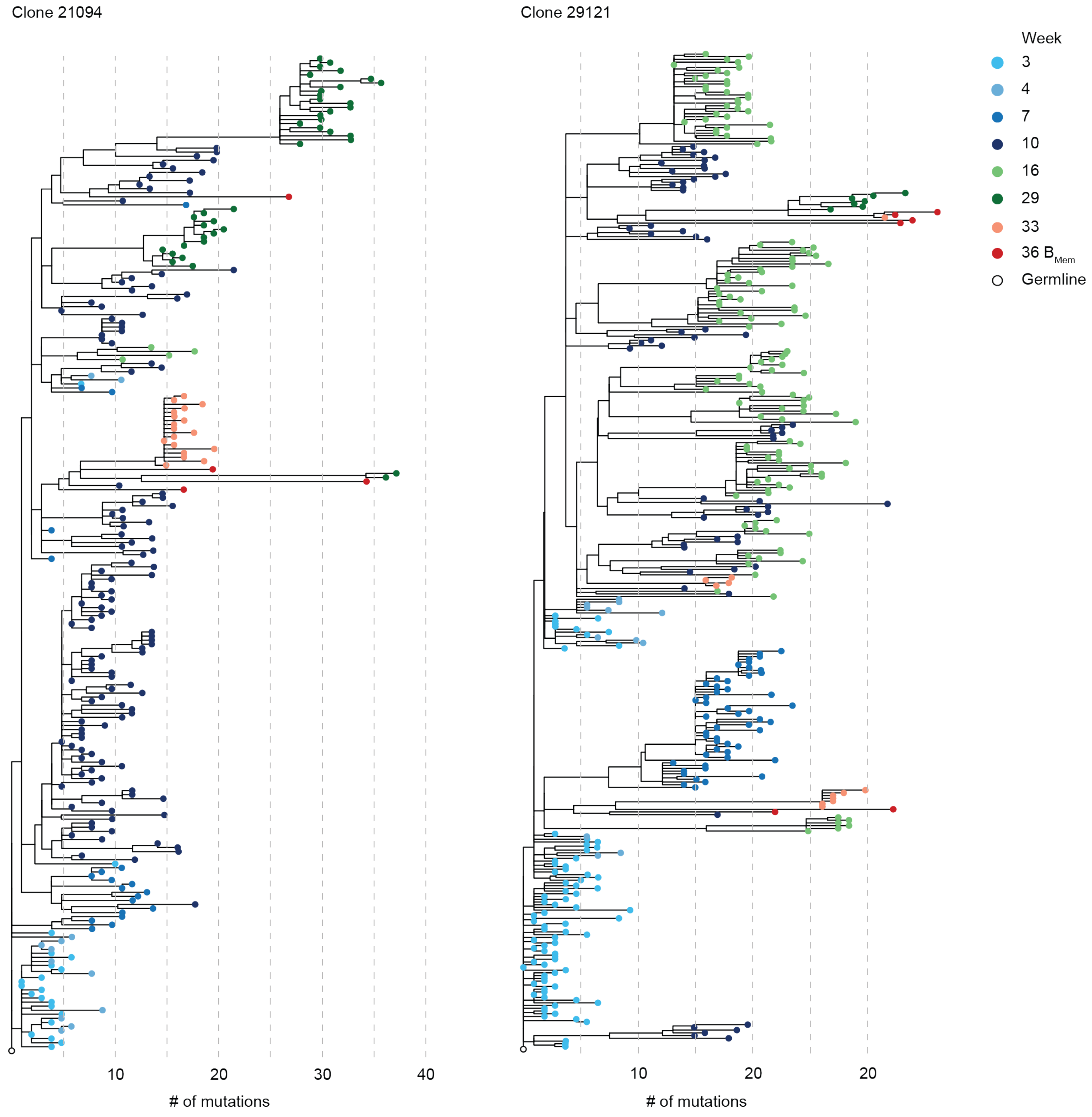


**Extended Data Fig 8. Linear examples of BCR clonal lineages.** Clonal families shown in Fig. 4k, observing clones over different LN FNA and PBMC sampling periods, represented as linear phylogenic trees.

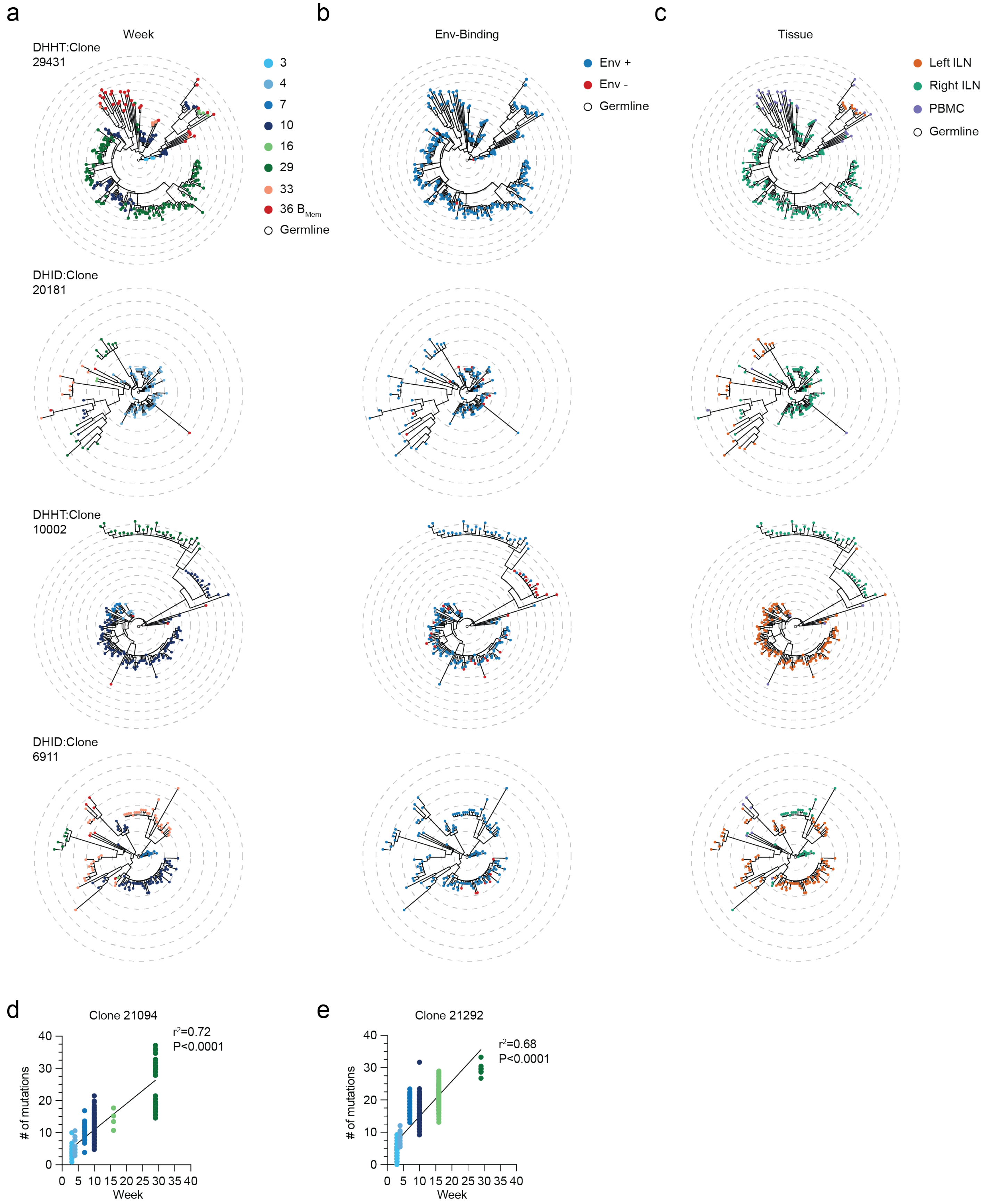


**Extended Data Fig 9. Additional examples of longitudinally assessed BCR clonal lineages. a,** Examples of clonal lineages observed over different LN FNA and PBMC sampling periods. **b,** B cells shown in (a) labeled by Env-binding based on flow cytometry. **c,** B cells shown in (a) labeled according to sampling location. Clones 29431, 20181 and 10002 all have H-CDR3s >14 AA in length, contained >2 N additions and >10 cells per LN. Clone 6911 has an H-CDR3 11 AA in length, contained >2 N additions and >10 cells per LN. Each ring indicates 5 HC mutations from the nearest common ancestor. **d-e,** Pearson correlation calculated between number of total HC mutations gained from the nearest common ancestor, and time post-prime. Correlation coefficients are calculated for clones 20194 (d) and 21292 (e) shown in Fig. 4k.

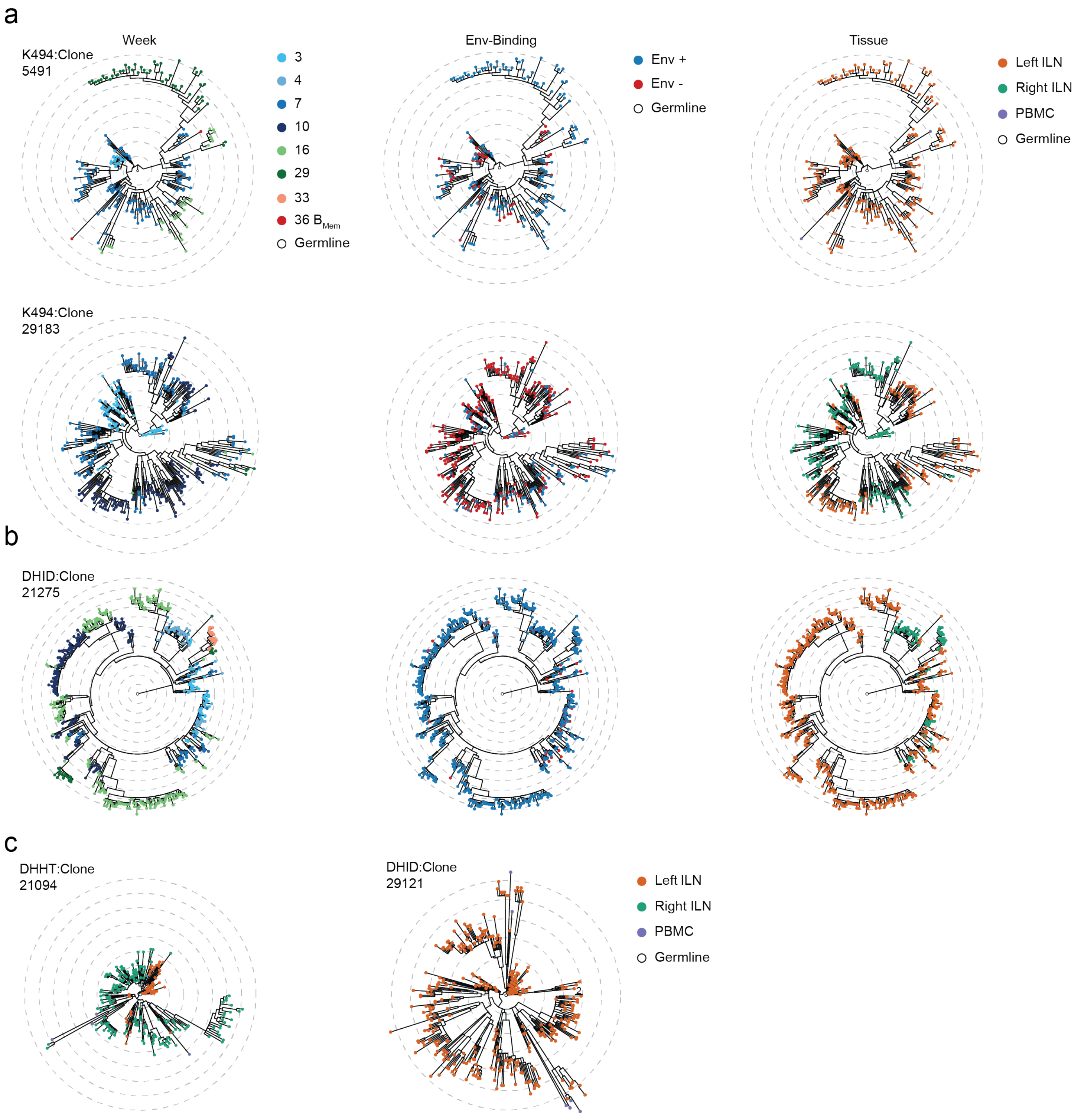


**Extended Data Fig 10. Examples of clonal lineages with unique features. a,** Clonal lineages where a substantial fraction of early $B_{GC}$ cells did not bind Env by flow cytometry. **b,** Example of a clonal lineage that was almost exclusively IgM. In (a) and (b), each B cell is labeled according to observed time point (left), Env-binding by flow cytometry (center), and sampling location (right). **c,** Lineage trees shown in Fig. 4k by sampling location. Clones 21275 and 20194 have H-CDR3s >14 AA in length, contained >2 N additions and >10 cells per LN.
